# Hyperglycemia promotes maladaptive Dectin-1 signaling and impairs skin antifungal host defense

**DOI:** 10.64898/2026.02.11.705431

**Authors:** Dante E. Reyna, Erin Davis, Ana C. G. Salina, Amondrea Blackman, Ruben Martinez-Barricarte, Amanda Doran, C. Henrique Serezani

## Abstract

People with chronic hyperglycemia are more susceptible to fungal skin infections, but the mechanisms underlying their worse clinical outcomes remain unclear. Using both *in vivo* and *in vitro* models, we explored how hyperglycemia influences skin antifungal defenses and how GLP1 agonists might restore host defense in diabetic conditions. Hyperglycemic mice showed increased susceptibility to *Candida albicans* skin infections, with larger lesions and higher fungal loads at all time points tested. Histology revealed larger abscesses, more extensive myeloid cell infiltration, and poorer control of fungal invasion, associated with increased chemoattractant production on day 1 post-infection. Despite heightened inflammatory responses, macrophages and keratinocytes exposed to high glucose exhibit markedly impaired fungal ingestion. RNAseq analysis of *C. albicans*-infected dermal macrophages cultured in high glucose showed enrichment of genes related to antimicrobial effectors and the C-type lectin receptor pathway, including *Clec7a* (Dectin-1), while suppressing downstream signaling pathways required for effective phagocytosis. Pharmacologic blockade or genetic deletion of Dectin-1 restored fungal uptake under high-glucose conditions and improved host defense *in vivo*. Mechanistically, Dectin-1 signaling in hyperglycemia promoted increased prostaglandin E₂ (PGE₂) production via induction of microsomal Prostaglandin E Synthase-1 (mPGES-1), and inhibition of PGE₂ synthesis rescued deficient phagocytic function. Finally, treatment with the glucagon-like peptide-1 (GLP-1) receptor agonist liraglutide reduced lesion size, fungal burden, inflammation, and tissue damage in diabetic mice, linking metabolic control to restoration of innate immune function. These findings identify maladaptive innate immune sensing as a key mechanism underlying susceptibility to fungal infection in diabetes and reveal how metabolic stress converts antifungal recognition pathways into drivers of inflammatory dysfunction.

## Introduction

Diabetes mellitus affects ∼ 537 million adults and is projected to rise to 783 million by 2045(1), with an estimated 6.7 million deaths attributable to diabetes or its related complications (2). Diabetes is associated with a markedly increased susceptibility to infections, particularly those affecting the skin and soft tissues (3). Among these, fungal infections caused by *Candida albicans* are disproportionately frequent, recurrent, and severe in individuals with chronic hyperglycemia. Cutaneous candidiasis represents a significant clinical burden, contributing to impaired wound healing, secondary bacterial infections, and increased hospitalization in diabetic patients (4). Despite this well-established association(5), the immunological mechanisms by which hyperglycemia compromises antifungal host defense in the skin remain incompletely understood. Recent Centers for Disease Control and Prevention (CDC) surveillance data indicate that approximately one-third of all candidemia cases in the United States occur in patients with diabetes (6).

Hyperglycemia disrupts multiple aspects of cellular physiology throughout the body, including immune cell function (7). Macrophages and monocytes from patients with diabetes exhibit decreased phagocytosis and reduced microbial killing compared with healthy controls (8). *In vivo*, skin infections in diabetic and/or obese mice result in exaggerated neutrophil migration and excessive production of inflammatory lipids and cytokines, leading to poor abscess formation and a failure to retain and clear pathogens in the skin (9). In diabetic wounds, macrophages struggle to transition from a pro-inflammatory phase to a healing state, thereby impairing healing and extending tissue damage (10,11).

Host defense against *C. albicans* involves rapid recognition by immune cells through pattern recognition receptors (PRRs) such as Dectin-1, the mannose receptor (MR), and toll-like receptors 2/4 (TLR2 and TLR4), which detect fungal components, including β-glucans and mannan (12–15). Dectin-1 specifically detects β-1,3-glucans during fungal growth, activating spleen tyrosine-protein kinase (SYK) and signaling through the CARD9–BCL10–MALT1 complex to activate the NF-κB and MAPK pathways (15,16). These pathways promote phagocytosis, pathogen destruction, and cytokine release. Dectin-1 also aids actin remodeling and phagocytic synapse formation, which are crucial for antifungal responses (17). Altered Dectin-1 signaling can weaken immunity, particularly in hyperglycemia and metabolic disorders, leading to persistent activation and ongoing inflammation associated with diabetes (18). In addition to recognizing microbial components, Dectin-1 can bind endogenous ligands, such as extracellular vimentin, and amplify inflammation, which can contribute to diabetes complications (19,20). It regulates sterile inflammation in diseases such as atherosclerosis, where activation increases cytokine production and leukocyte recruitment, often disrupted in metabolic diseases (20). Understanding the role of Dectin-1 in inflammation is vital to understanding its impact on tissue-specific inflammation and infection in diabetes.

Among the mediators produced by macrophages following β-glucan stimulation and Dectin-1 signaling, one is prostaglandin E₂ (PGE₂) (21). PGE₂ is synthesized from arachidonic acid via cyclooxygenase (COX) and mPGES1-2 enzymes and acts as a lipid mediator at inflammatory sites, with both pro- and anti-inflammatory effects depending on receptor interactions (22,23). It functions as a vasodilator, increasing endothelial permeability and cell recruitment, but also suppresses inflammatory cytokine production and reduces microbial killing by lowering ROS levels (24). Our previous work showed an immune imbalance in the skin of infected hyperglycemic/obese mice, with an early rise in LTB_4_ and a decrease in PGE₂ from day 3 (9). Topical misoprostol restored PGE₂ and reduced the bacterial load in the skin of hyperglycemic and obese mice (25). These findings suggest that alterations in PGE₂ signaling play a crucial role in linking metabolic disorders, inflammation, and susceptibility to infection.

Despite immune defects linked to hyperglycemia that can increase infection risk, the specific immune interactions and pathogenic processes that increase susceptibility to *C. albicans* in individuals with diabetes remain poorly understood. Our study reveals that hyperglycemia aggravates infection severity, as evidenced by larger lesions, higher fungal burdens, and greater production of inflammatory mediators, which lead to tissue damage. Importantly, we identify a novel role for Dectin-1 in inhibiting *C. albicans* ingestion, suggesting that its function under metabolic stress differs from its traditional role in antifungal defense. These results highlight how hyperglycemia impairs antifungal effector functions and provide new insights into why diabetes predispose individuals to fungal infections. Furthermore, we found that treating diabetic mice with a GLP-1 agonist reduces lesion size and fungal burden, suggesting potential avenues for new therapies.

## Materials and Methods

### Animals

C57BL/6J breeding pairs were obtained from The Jackson Laboratory (Bar Harbor, ME, USA), and colonies were maintained at Vanderbilt University Medical Center (VUMC) in Nashville, TN, USA. Male mice aged 6–12 weeks were used for all experiments. Animals were housed with ad libitum access to food and water. All procedures were approved by the Vanderbilt University Medical Center Institutional Animal Care and Use Committee (IACUC) and conducted in accordance with the NIH Guidelines for the Care and Use of Laboratory Animals.

### Isolation of skin resident macrophages

Mice were shaved, and fur was removed with a depilatory agent. Dorsal skin was excised and digested with collagenase IV (Merck KGaA, Darmstadt, Germany), as described (25). The resulting cell suspension was stained and sorted to isolate tissue macrophages (F4/80^+^ CD68^+^ double-positive cells). The isolated macrophages were resuspended in complete Dulbecco’s Modified Eagle’s Medium (DMEM) supplemented with 10% heat-inactivated fetal bovine serum (FBS), 2 mM L-glutamine, 100 units/mL penicillin, 100 μg/mL streptomycin, 25 mM HEPES, and 2.5 × 10^5^ M 2-mercaptoethanol (all from Gibco Invitrogen, Grand Island, NY, USA). They were then seeded into culture plates and incubated with 100 ng/mL murine M-CSF (Sigma-Aldrich, St Louis, MO, USA).

### Immortalization of macrophages

Dermal macrophages (DMs) and bone marrow–derived macrophages (BMDMs) from C57BL/6 and Dectin-1–deficient mice were immortalized using the J2 retroviral system. Primary DMs were isolated as previously described (26). DMs were isolated using a macrophage isolation kit (Myltenyl) and seeded at 3 × 10⁵ cells/mL in 12-well plates (Costar, Cambridge, MA) and cultured in DMEM containing 100 ng/mL M-CSF for 24 h to promote proliferation. DMCLs were generated and prepared for immortalization as previously reported (26).

To immortalize bone marrow-derived macrophages (iBMDMs), bone marrow cells from C57BL/6 and Dectin-1 knockout (KO) mice were seeded at 1x 10^6^ cells/mL and cultured in 10 mL DMEM containing 20 ng/mL M-CSF for 48 hours. On day 3, cells were infected with J2 virus supernatants mixed with DMEM containing 20 ng/mL M-CSF, and polybrene was added to the plates. Immortalization was performed as above.

### Cell culture

All immortalized macrophages were maintained in culture in DMEM supplemented with 10% heat-inactivated FBS and 1% penicillin-streptomycin at 37°C in 5% CO₂. To simulate different glucose concentrations, cells were incubated in media containing either 5 mM (low glucose) or 25 mM (high glucose) D-glucose for at least 24 hours before infection or stimulation.

### Human keratinocytes culture

Immortalized human keratinocytes (N/TERT) were cultured in Keratinocyte Serum-Free Medium (K-SFM) (Thermo Fisher Scientific), supplemented with pituitary gland extract, epidermal growth factor, and 0.4 mM CaCl₂ to support cell adhesion and monolayer stability, and maintained at 37 °C in 5% CO₂ (27). Cells were passaged at approximately 50% confluence for routine culture and allowed to reach full confluence before experimental assays.

To model hyperglycemic conditions, D-glucose was added to the basal low-glucose (5 mM) K-SFM to a final concentration of 25 mM, and the mixture was filter-sterilized. Cells were equilibrated in the indicated glucose conditions before downstream experimental manipulation, as described below.

### *C. albicans* strains and culture preparation

The *C. albicans* CAF2.1 tdTomato strain, a URA3-reconstituted derivative of SC5314 that retains wild-type virulence and filamentation, was provided by Mikhail Lionakis (National Institutes of Health, Bethesda, MD) (28,29). The tdTomato construct was integrated using a NAT1 cassette under a constitutive promoter to enable fluorescent visualization during infection (28).

*C. albicans* was streaked onto Sabouraud Dextrose Agar (BD Difco) to obtain isolated colonies and incubated at 30°C for 48 hours. A single colony was inoculated into 30 mL of Sabouraud Dextrose Broth (BD Difco) and cultured at 30°C with shaking (200 rpm) for 48 hours. Yeast cells were counted using a hemocytometer with 0.4% trypan blue exclusion to verify cell density and viability. Cultures were pelleted at 3,000 × g for 5 minutes, resuspended in sterile 20% glycerol, aliquoted, and stored at -80°C. A fresh aliquot was thawed, washed once in PBS, and used for each experimental infection to ensure consistency across experiments.

A clinical commensal *C. albicans* isolate, strain 838-4 was a gift from Dr O’Meara (University of Michigan). This strain constitutively expresses an infrared fluorescent protein (iRFP) and was used as a representative colonization-associated strain, as previously described (30).

Strain 838-4 was cultured, expanded, and cryopreserved under the same conditions as CAF2-1 tdTomato to ensure experimental consistency across infection models

### Induction of *C. albicans* hyphal growth

To induce hyphal morphogenesis, *C. albicans* yeast cells were cultured under serum-containing conditions at 37°C. *C. albicans* SC5314 aliquots were thawed and cultured overnight in Sabouraud Dextrose Broth (BD Difco), washed in PBS, and resuspended in fresh broth supplemented with 10% FBS at 37°C for 2 hours to induce hyphal formation (31). Following incubation, cells were harvested and processed for downstream applications as described below.

### Induction of hyperglycemia

Male mice aged 6–8 weeks were injected with streptozotocin (STZ) to induce hyperglycemia. Mice received intraperitoneal injections of STZ (50 mg/kg body weight; Sigma-Aldrich) once daily for 5 consecutive days (32). STZ was freshly prepared each day in a cold 0.1 M citrate buffer (pH 4.5) and administered within 15 minutes of preparation. Control animals received citrate buffer alone. Blood glucose levels were measured starting 7 days after the final injection using a handheld glucometer (Accu-Chek). Mice with non-fasting blood glucose levels exceeding 250 mg/dL were classified as hyperglycemic. All experiments were performed 30 days after initiation of STZ injections.

### *Candida albicans* skin infection model

Male C57BL/6 mice aged 8 to 12 weeks, treated with STZ or vehicle, were used. Mice were infected subcutaneously (s.c.) at two dorsal sites with approximately 1 × 10⁷ *C. albicans* cells (50 µL per site) prepared from frozen aliquots (33). Lesion size was measured with digital calipers every other day, and the affected area was calculated as length × width (mm²). At specific time points (days 1, 3, and 7 post-infection), skin biopsies were taken using an 8-mm punch after CO₂ euthanasia and ethanol sterilization of the skin.

For Liraglutide (Novo Nordisk, Bagsvaerd, Denmark) experiments, mice received s.c. Liraglutide at 0.05 mg/kg once daily, as described (34). Liraglutide was prepared in 0.05% BSA–PBS, which also served as the vehicle control. Treatment began two days before the infection. Liraglutide dosing was adjusted based on mouse weight at each injection. After two days of pretreatment and glucose monitoring, mice were infected with 1 × 10⁷ *C. albicans* and continued to receive daily injections of Liraglutide or vehicle, along with glucose monitoring, until day 7 post-infection. Blood glucose levels were measured and recorded as described.

### Laminarin-mediated Dectin-1 inhibition

Diabetic and non-diabetic mice received a s.c. injection of 250 mg/kg laminarin into the middle of their backs, near the infection site, for 6 hours before *C. albicans* injection. At 24 hours post-infection, skin biopsies were collected and processed for downstream analyses.

### Fungal burden quantification

Skin biopsies were collected, weighed, and homogenized in 300 µL of Sabouraud dextrose broth. Serial dilutions of the homogenates were prepared, and 10 µL of each dilution was plated onto Sabouraud dextrose agar. Plates were incubated overnight at 37 °C, and colony-forming units (CFUs) were enumerated and adjusted for tissue weight. Results are presented as CFUs per milligram of tissue.

### Histology

Biopsies were fixed overnight in 10% formalin at 4°C and subsequently processed for 18 hours following paraffin embedding. Sections were stained with hematoxylin & eosin (H&E), Masson’s Trichrome, or Periodic Acid Schiff (PAS), as we have done before (35). Slides were imaged with a Cytation 7 Cell Imaging Multi-Mode Reader (BioTek) at 20× magnification. All histological images shown are montages generated by stitching multiple adjacent 20× fields to ensure comprehensive visualization of lesion architecture.

### RNA isolation and qRT-PCR

Samples were collected at various time points after infection and homogenized in RLT buffer (QIAGEN, Germantown, MD). Total RNA was extracted using the RNeasy Mini Kit (QIAGEN) according to the manufacturer’s instructions. One microgram of RNA was reverse-transcribed into cDNA using the iScript cDNA Synthesis Kit (Bio-Rad, Hercules, CA). Quantitative PCR (qPCR) was performed on a CFX96 Real-Time PCR Detection System (Bio-Rad) with Platinum SYBR Green Master Mix (Invitrogen, Carlsbad, CA). Gene expression levels were normalized to β-actin and calculated using the comparative threshold cycle (ΔΔCt) method. Primer sequences for β-actin and target genes were obtained from Integrated DNA Technologies (IDT, Coralville, IA).

### RNA-seq Processing and Functional Pathway Enrichment Analysis

Dermal macrophage cell line (DMCL) was cultured under low-glucose (5 mM) or high-glucose (25 mM) conditions for 24 hours and was either challenged with *C. albicans* (multiplicity of infection [MOI] 5) for 18 hours or left unchallenged. RNA was harvested using the RNeasy Mini Kit (Qiagen, #74104). Purified RNA was submitted for processing at VANTAGE (Vanderbilt Technologies for Advanced Genomics), as we have previously shown (36). Genes with log_2_ FoldChange (log_2_FC) values greater than 2 or less than –2 and *P* values <0.05 were used to generate log-normalized volcano plots. Differential expression analysis was performed on raw counts using DESeq2. Genes with adjusted P values below 0.05 and an absolute fold change of at least 1 were classified as significantly differentially expressed. Upregulated and downregulated gene sets were separated, and log-normalized data were used for visualization.

Functional enrichment analysis was conducted using the clusterProfiler suite in R (37). Enriched Gene Ontology Biological Process terms, KEGG pathways, and Reactome signaling pathways were identified based on ENTREZ-mapped differentially expressed genes (34–36). Analyses were performed with a Benjamini–Hochberg-adjusted P-value cutoff of 0.05, using the complete set of detected genes as the background. Results were visualized through fold enrichment dot plots, KEGG enrichment maps, and KEGG gene–pathway connectivity networks colored by fold change.

### Phagocytosis assay

WT and Dectin-1 KO iBMDM and DMCL were cultured in low- or high-glucose media for at least 24 hours, then challenged with CAF2-1 tdTomato C. *albicans* at an MOI of 5:1 (yeast: macrophage). Where indicated, laminarin (100 µg/mL) or the mPGES1 inhibitor PF-03549184 (1 μM) was added 20 minutes before infection. After 2 hours of incubation, cells were washed with PBS, extracellular fluorescence was quenched with 0.2% trypan blue, and intracellular fluorescence was quantified using a plate reader. All experiments were performed in technical triplicate and repeated at least three times independently, using a 96-well black plate (Corning).

### Western blotting

Western blots were conducted as previously described (41). Protein samples were subjected to SDS-PAGE, transferred to nitrocellulose membranes, and probed with primary antibodies: mPGES-1 (Cell Signaling Technology). After washing, the membranes were incubated with suitable fluorophore-conjugated secondary antibodies (1:10,000, anti-rabbit IgG, IRDye 800CW, #926-32211, Licor). Band intensities were quantified using ImageJ software (NIH), following the same procedure as in (42).

### Multiplex bead array

Biopsy samples were collected, weighed, and homogenized in 200 μL of TNE cell lysis buffer containing phosphatase and protease inhibitors. The samples were then centrifuged to remove cellular debris. Skin biopsy homogenates were analyzed using the pro-inflammatory 18-plex Discovery Assay from Eve Technologies (Calgary, AB) to measure cytokines and chemokines (43). All concentrations were normalized to the total protein content of the corresponding tissue lysates and expressed as ng/mg protein.

### Detection of PGE_2_, IL-1β and IL-10

PGE₂ in the skin: Skin biopsies were collected, weighed, and rapidly frozen in liquid nitrogen. PGE₂ abundance was determined using EIA, according to the manufacturer’s instructions (Cayman Chemical, Ann Arbor, MI).

For IL-1β and IL-10 detection, skin biopsies were homogenized in 200 μL of TNE cell lysis buffer supplemented with protease and phosphatase inhibitors. Cytokine levels, including IL-10/IL-1β, were determined using BioLegend ELISA kits according to the manufacturer’s instructions (BioLegend, San Diego, CA). All analyte concentrations were normalized to total protein content in the corresponding tissue lysates and are expressed as pg/mg protein.

### Semiquantitative phospho-array

DMCL were cultured in low- or high-glucose media and challenged with *C. albicans* for 22 hours. Protein content was quantified by Bradford assay, and 50 μg of protein was used for qualitative measurement of phosphorylated proteins using the Proteome Profiler Human Phospho-Kinase Array (ARY003C), as recommended by the manufacturer (R&D Systems, Wiesbaden, Germany).

### Statistical Analysis

All statistical analyses were performed using GraphPad Prism (GraphPad Software, San Diego, CA). One-way ANOVA with Tukey’s multiple comparisons test was used to compare three or more experimental groups. Data are presented as mean ± SEM. For experiments involving two independent variables, statistical significance was determined using two-way ANOVA with Sidak’s multiple-comparisons test to correct for multiple testing across two families of comparisons. For two-group comparisons in which data did not meet the assumptions of normality, a Mann–Whitney U test was used. A p-value < 0.05 was considered statistically significant.

## Results

### Hyperglycemia exacerbates cutaneous *C. albicans* infection, leading to a greater fungal burden and tissue pathology

To assess the effect of hyperglycemia on cutaneous *C. albicans* infection, we developed a mouse model of diabetic skin candidiasis using STZ-induced diabetes and non-diabetic control mice over 30 days. Diabetic mice developed significantly larger and longer-lasting lesions than controls during the 7-day infection period (mean lesion area of 72 ± 0.5476 mm² in diabetics versus 62 ± 0.5476 mm² in controls at day 4, p < 0001) (Figure 1A). We demonstrated that larger lesion size correlates with impaired fungal clearance. Fungal burden was measured in skin biopsies collected on days 1, 3, and 7. Our data show that although non-diabetic mice showed reduced CFU over time, infected diabetic mice had notably higher CFU counts across all time points (Figure 1B). Next, we sought to determine whether diabetes also predisposes to other *C. albicans* strains with reduced hyphal formation (the invasive form of the disease). Our data show that infection with the commensal isolate 838-4 also resulted in increased lesion size and fungal burden in diabetic mice compared with controls (Supplementary Figure 1). Together, these findings indicate that hyperglycemia impairs fungal control early during infection, independent of hyphal induction, consistent with a disruption of initial innate immune responses.

**Figure 1.**
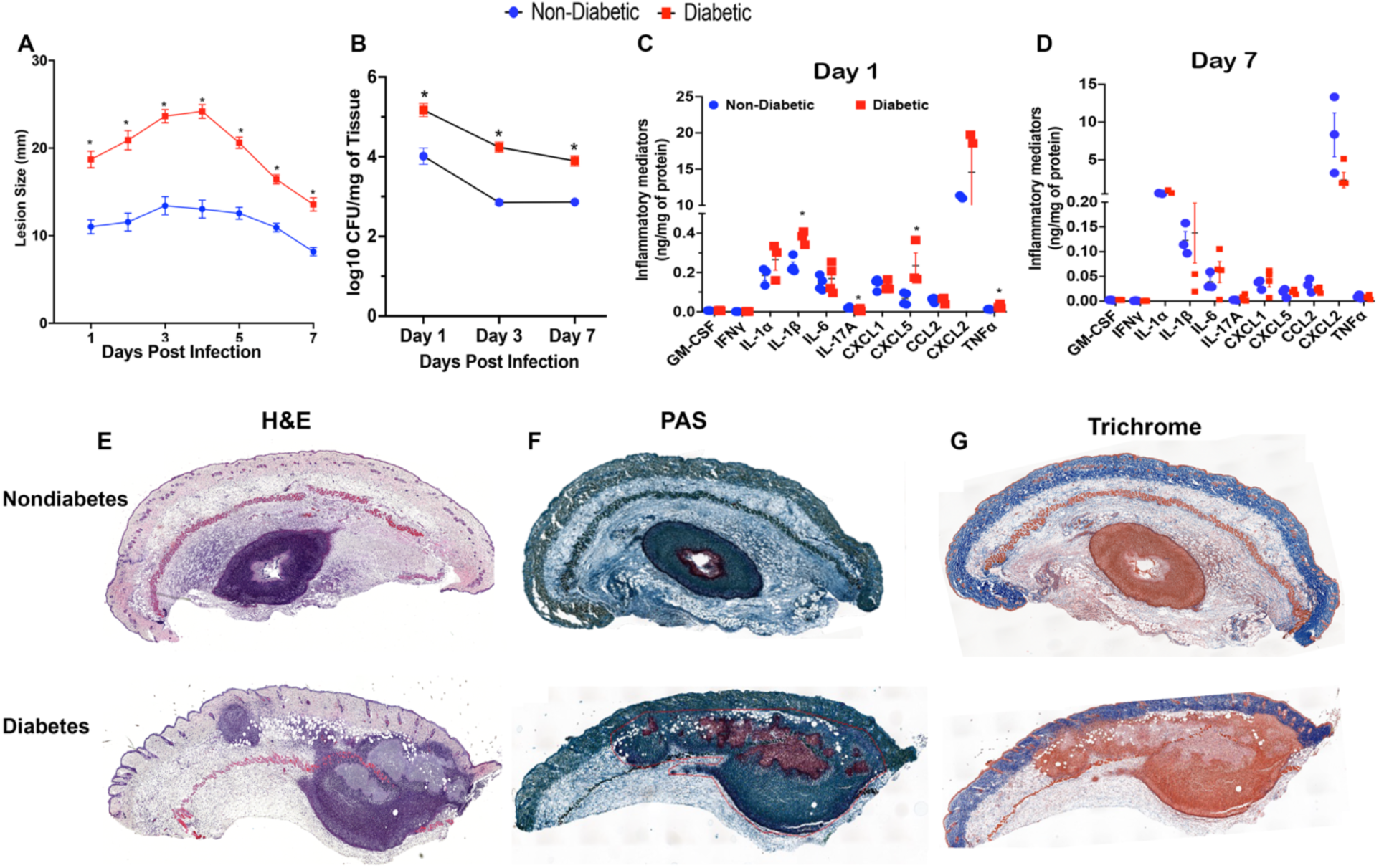
Diabetes predisposes mice to *Candida albicans* skin infection. A) Diabetic and non-diabetic C57BL/6 mice were infected s.c. with 1×10^7^ *C. albicans* yeast, and lesion size was monitored daily for 7 days. B) CFU of *C. albicans* in diabetic and non-diabetic mice on days 1, 3, and 7 post-infection. C-D) Skin sections from diabetic and non-diabetic mice were collected on days 1 (C) and 7 (D) post-infection and subjected to cytokine bead array. E) Sections were stained with H&E, F) Periodic-Acid Schiff (PAS), and G) Masson trichrome blue. In all cases, at least 4 mice/group were analyzed. Data represent mean ± SEM. *p < 0.05 vs non-diabetic control by Mann–Whitney test (N = 4 controls, 7 diabetics).

Next, we characterized the inflammatory milieu in infected skin from diabetic and non-diabetic mice. We first performed cytokine bead array analysis of skin lysates on days 1 and 7 post-infection. On day 1, only IL-1β, IL-17A, TNF, and CXCL5 levels are increased in the infected skin of diabetic mice (Figure 1C). On day 7, we did not detect differences in the abundance of the cytokines tested (Figure 1D), suggesting that early responses dictate the outcome of *C. albicans* infection in diabetic mice.

We then analyzed overall tissue pathology in infected diabetic and non-diabetic mice. H&E staining revealed differences in tissue structure and inflammatory responses between the two groups after *C. albicans* infection. In non-diabetic mice examined 24 hours after infection (top panel), skin architecture was mostly intact, with the epidermis and dermis preserved. A well-defined, localized inflammatory focus was observed in the dermis and subcutaneous tissue, characterized by an accumulation of inflammatory cells. The lesion center showed early necrosis and cellular debris (Figure 1E). Conversely, diabetic mice (bottom panel) exhibited significant disruption of normal tissue structure. Extensive inflammatory infiltrates were observed throughout the dermis and epidermis, along with interstitial edema and dilated blood vessels. Large areas of tissue necrosis and debris were present, with poorly defined lesion boundaries, indicating a poorly contained inflammatory response (Figure 1E). In non-diabetic skin infections, we observed smaller inflammatory areas, less dermal thickening, and lower necrosis rates than in diabetic skin infections. These data reveal that diabetes alters early inflammatory skin responses to *C. albicans*, leading to a more severe inflammatory response and greater tissue damage at 24 hours post-infection.

PAS staining revealed a more widespread and denser fungal presence in diabetic lesions (Figure 1F). Masson’s trichrome staining revealed clear differences in skin collagen structure between the two tested groups of animals following *C. albicans* infection. In infected non-diabetic mice (top panel), dermal collagen fibers remained well-organized and preserved, with densely packed collagen localized around the inflammatory site. In the infected skin of diabetic mice (bottom panel), we observed disruption and loss of dermal collagen, characterized by collagen-depleted regions that coincided with areas of intense inflammation and tissue damage. Together, these results establish a model of diabetic candidiasis and show that hyperglycemia exacerbates lesion severity, fungal burden, and abscess development *in vivo*. This model provides a framework for exploring how diabetes impairs antifungal immune responses amid metabolic disturbances.

### The GLP1 agonist Liraglutide reduces tissue damage severity in infected diabetic mice

Because metabolic dysregulation worsens immune dysfunction and inflammation in diabetes (44), we next investigated whether improving metabolic status could modify the severity of infection. We treated animals daily with the GLP-1 receptor agonist liraglutide, a clinically approved therapy for glycemic control in patients with diabetes (45), throughout the 7-day infection. Liraglutide significantly reduced circulating glucose levels in diabetic mice, with non-fasting blood glucose levels normalizing to those of non-diabetic controls by day 1 of treatment and remaining controlled during continued administration (Figure 2A). While diabetic mice treated with vehicle developed significantly larger, more ulcerated lesions than all other groups, liraglutide treatment markedly reduced lesion size in diabetic mice (Figure 2B). Decreased lesion size, correlated with reduced fungal load in diabetic mice treated with liraglutide (Figure 2C). H&E staining showed that liraglutide-treated diabetic mice had decreased inflammatory infiltrate and abscesses (Figure 2D). Fungal stain (PAS) showed higher fungal loads in vehicle-treated diabetic mice than in liraglutide-treated diabetic mice (Figure 2D). Reduced lesion severity and tissue damage suggest alterations in the skin’s inflammatory milieu in these animals. To test this hypothesis, we measured cytokines involved in phagocyte function, e.g., IL-1β and IL-10, in whole-tissue lysates.

**Figure 2.**
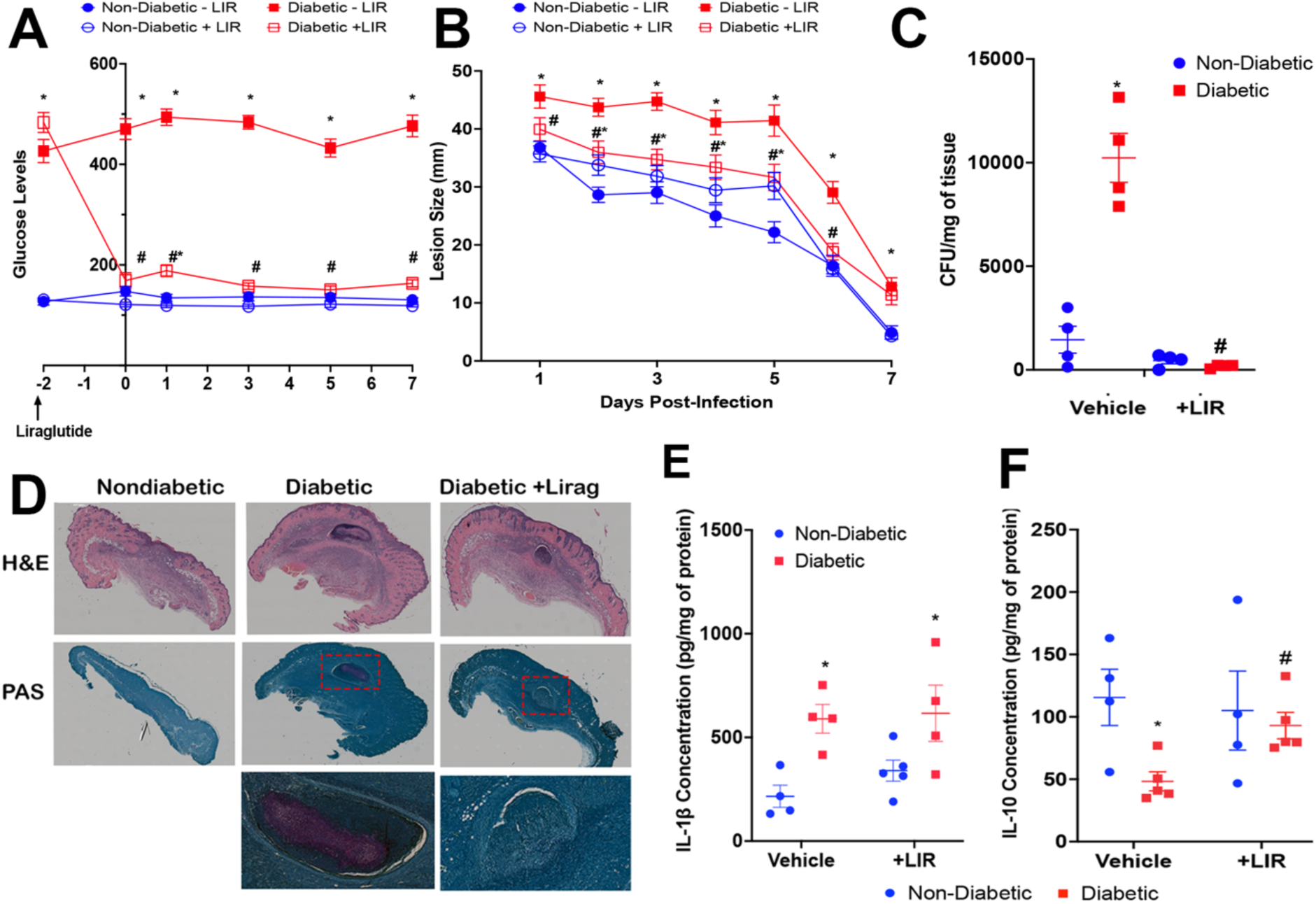
Liraglutide treatment improves host defense against *C. albicans infection in* diabetic mice. Diabetic and non-diabetic mice were treated with or without Liraglutide, followed by *C. albicans* skin infection. **A)** Glucose levels were determined every other day before and after *C. albicans* infection. **B)** Lesion size was measured every other day for 7 days. **C)** CFU determination in the infected skin of diabetic and nondiabetic mice treated or not with liraglutide. **D)** Skin biopsies were collected on day 7 post-infection, and skin sections were stained with H&E and PAS. Red boxes indicate fungal detection. **E-F)** Detection of IL1β **(E)** and IL10 **(F)** in skin lysates from liraglutide-treated diabetic mice. Data represent mean ± SEM. *p < 0.05 vs non-diabetic control and # p < 0.05 vs diabetic mice by Mann–Whitney test. n = at least 4/group

Although IL-1β was increased in diabetic mice, we did not observe any changes in IL-1β among the liraglutide-treated groups (Figure 2E). Our data showed that *C. albicans* decreases IL-10 in diabetic mice, and that liraglutide restores IL-10 to levels observed in non-diabetic, infected mice (Figure 2F). These data suggest that liraglutide restores glucose levels in diabetic mice, thereby inducing a balanced inflammatory response that reduces tissue damage and improves microbial clearance.

### Hyperglycemia impairs fungal ingestion in different cell types

One of the earliest steps in infection control is the recognition and phagocytosis of pathogens by innate immune cells such as macrophages. To determine whether hyperglycemia alters macrophage fungal ingestion, we compared the uptake of *C. albicans* and zymosan particles across multiple macrophage types and glucose levels.(5) We first measured fungal ingestion in peritoneal macrophages (PMs) from diabetic and non-diabetic mice. Our data show that PMs from diabetic mice exhibit reduced phagocytic capacity compared with those from non-diabetic mice (Figure 3A). Next, we assessed whether high glucose impairs ingestion in dermal macrophages. DMCLs cultured in high-glucose media showed significantly lower phagocytosis than those cultured in low-glucose media (Figure 3B, 3C, and 3E) for both *C. albicans* yeast and zymosan (Figure 3B, 3C, and 3E). The same pattern was observed in immortalized bone marrow-derived macrophages (iBMDM), indicating a broad inhibitory effect of high glucose on fungal ingestion (Figure 3D). Next, we sought to determine whether high glucose inhibits the ingestion of different fungal forms (Figure 3E). Our data show that high glucose decreases phagocytosis of both yeast and hyphal forms by macrophages.

**Figure 3.**
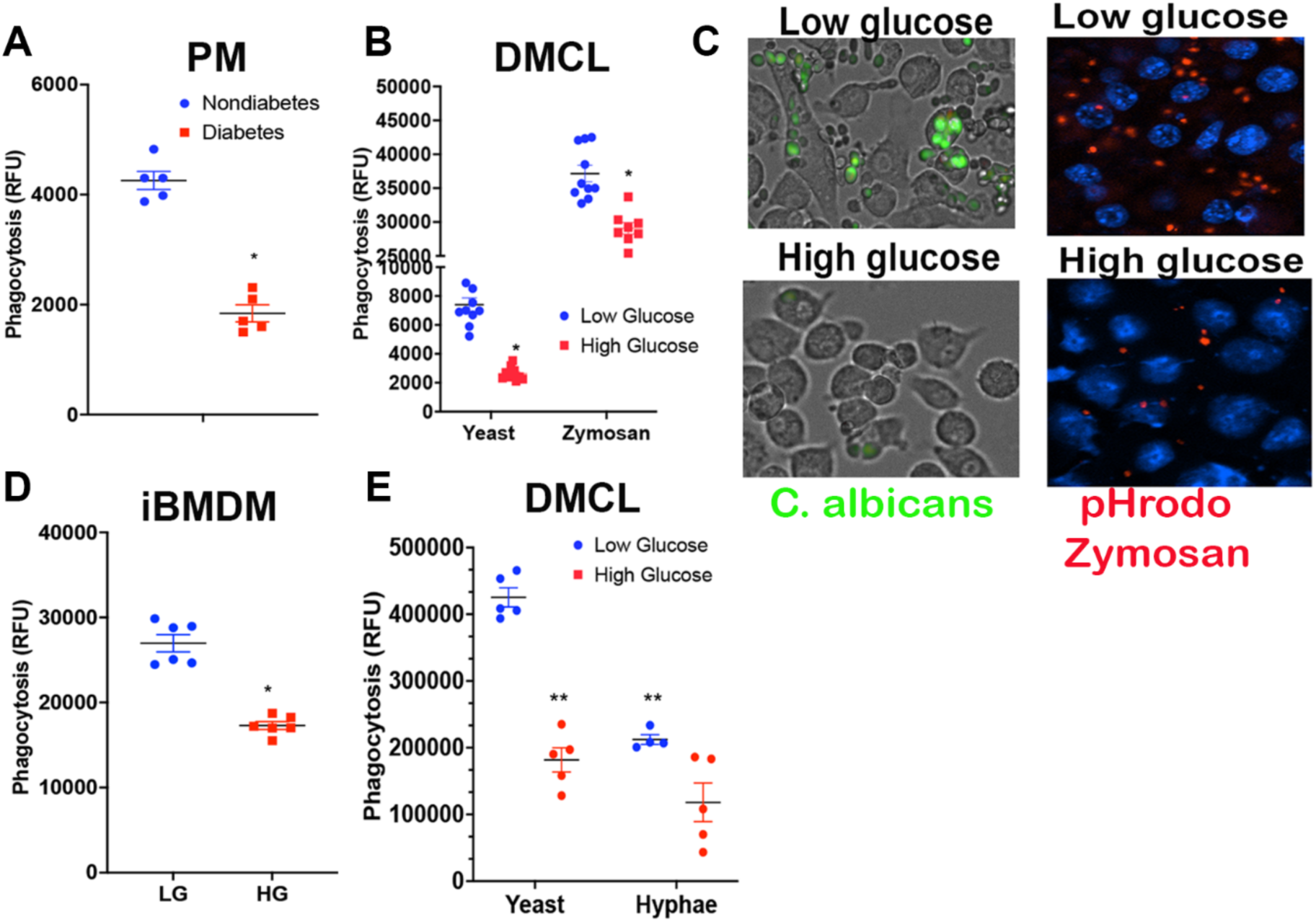
High glucose inhibits *C. albicans* ingestion across different macrophage populations. **A)** Peritoneal macrophages (PM) from diabetic and non-diabetic mice were challenged with an MOI of 5 *C. albicans* for 1 hour, followed by determination of phagocytosis. **B)** DMCLs were cultured in low (5 mM) or high (25 mM) glucose media for at least 48 hours, then challenged with an MOI of 5 *C. albicans* yeast or pHrodo-labeled zymosan. **C)** DMCLs were cultured and infected as in B and imaged to detect intracellular *C. albicans* **(left)** or pHrodo-zymosan **(right)**. **D)** iBMDM were cultured as in B, and phagocytosis was determined. **E)** DMCLs were cultured in low or high glucose and challenged with *C. albicans* yeast or hyphae at an MOI of 5, and phagocytosis was determined as described in the methods. Data represent mean ± SEM. *p < 0.05 vs non-diabetic control or low glucose media by Mann–Whitney test. n = at least 4/group.

### RNA-seq analysis identifies glucose-dependent remodeling of macrophage antifungal pathways

Dermal macrophages orchestrate both the inflammatory response and fungal clearance (46), so we aimed to define the transcriptional programs selectively altered by high glucose at the cellular level. To do so, we performed RNA-seq analysis on *C. albicans-*challenged DMCL cultured under low- or high-glucose conditions. We detected 1585 genes upregulated and 963 downregulated in cells cultured in high glucose (Figure 4A). KEGG analysis showed that genes involved in phagosome formation and NF-κB signaling were downregulated. Increased gene enrichment was observed for cytokine-cytokine receptor interaction and the cytoskeleton (Supplementary Figure 3). Next, we generated a heat map of gene clusters (Supplementary Figure 3A). KEGG analysis identified 3 significant clusters (Supplementary Figure 3B). Cluster 3 contains genes involved in *Staphylococcus aureus* infection, cell adhesion, and hematopoietic cell lineage. Cluster 4 shows enrichment in genes related to cell metabolism. Cluster 5 is restricted to neuroactive ligand-receptor interaction. GO biological function analysis showed that cluster 3 is enriched for genes involved in host defense and cell motility. Cluster 4 comprises genes that regulate cell metabolism and leukocyte function (Suppl. Fig 3B). Further examination of CLR genes demonstrates increased expression of dectin-1 and -2 and decreased expression of several downstream components of dectin signaling, including *Syk, Malt1,* and the p38 MAPKs *Mapk13 and Mapk14*.

**Figure 4.**
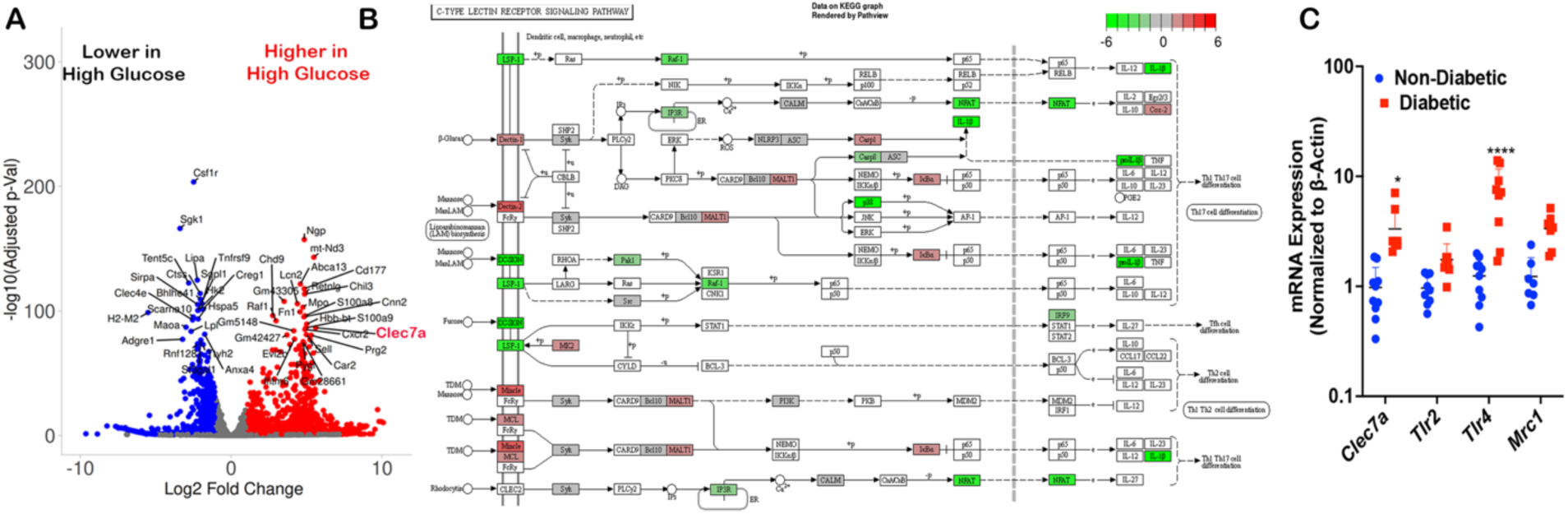
High glucose increases the expression of genes involved in pathogen recognition in *C. albicans*-infected dermal macrophages. **A-D)** DMCLs were cultured in low- or high-glucose conditions and challenged with *C. albicans* for 12 hours. **(A)** Volcano plot (log-transformed) shows significantly up-or downregulated genes. The genes listed are involved in macrophage antimicrobial effector function and inflammatory responses. The iDEP Analysis Toolkit and FDR < 0.05 were used to perform **(B)** KEGG map of differentially expressed genes involved in C-type lectin receptor signaling pathways. **C)** Expression of the indicated genes in the skin 24 hours after *C. albicans* infection in diabetic and non-diabetic mice, determined by qPCR as described in Materials and Methods. Data represent at least 3 independent experiments. Data represent mean ± SEM. *p < 0.05 vs non-diabetic control by Mann–Whitney test. n = at least 5/group

To better comprehend how these pathways interact, we created a gene connectivity network based on the KEGG-enriched gene set (Figure 3B). This network revealed a connected cluster centered on *Syk*, *Malt1*, *Ptgs2*, and *Il1b*, suggesting coordinated inhibition of the Syk–Malt1 axis and inflammatory effector genes associated with increased prostaglandin production and NF-κB activation (Figure 3B). Next, we examined the expression of receptors that could recognize *C. albicans* in the skin of diabetic mice. Interestingly, we found increased levels of *Clec7a* (dectin-1) and *Tlr4*, but not *Tlr2* or *Mrc1* (mannose receptor) (Figure 3C). These findings imply that hyperglycemia selectively enhances *Clec7a-*and *Tlr4*-related pathways rather than globally boosting innate receptor expression. Since high glucose specifically diminishes fungal recognition and inflammatory gene programs, including CLR signaling, NF-κB activation, and prostaglandin synthesis, we then assessed whether this transcriptional downregulation impairs macrophage recognition and fungal phagocytosis.

### Dectin-1 inhibits phagocytosis under high-glucose conditions

The decreased uptake of the β-glucan particle zymosan in high glucose, along with Dectin-1’s role as the main receptor for β-glucan phagocytosis and the observed increase in Dectin-1 expression in macrophages cultured in high glucose, indicates that hyperglycemia might interfere with Dectin-1-dependent processes engulfment. To test this hypothesis, we initially treated DMCLs cultured in low- and high-glucose media with the Dectin-1 antagonist laminarin, then challenged the cells with *C. albicans*. Our data show that laminarin decreases fungal ingestion in DMCLs cultured in low-glucose media and greatly increases phagocytosis in DMCLs cultured in high-glucose media (Figure 5A). To build on this observation, we cultured and challenged WT and Dectin-1 KO iBMDMs under low- and high-glucose conditions and found that WT iBMDMs exposed to high glucose exhibited lower phagocytic activity than WT cells cultured in low glucose, independent of the MOI tested (Figure 5B). In contrast, Dectin-1 KO iBMDMs showed higher phagocytic activity and maintained higher uptake (Figure 5B) only when cells were maintained in high glucose. These data indicate that the phagocytic defect induced by hyperglycemia is specific to WT iBMDMs and dependent on Dectin-1 signaling.

**Figure 5.**
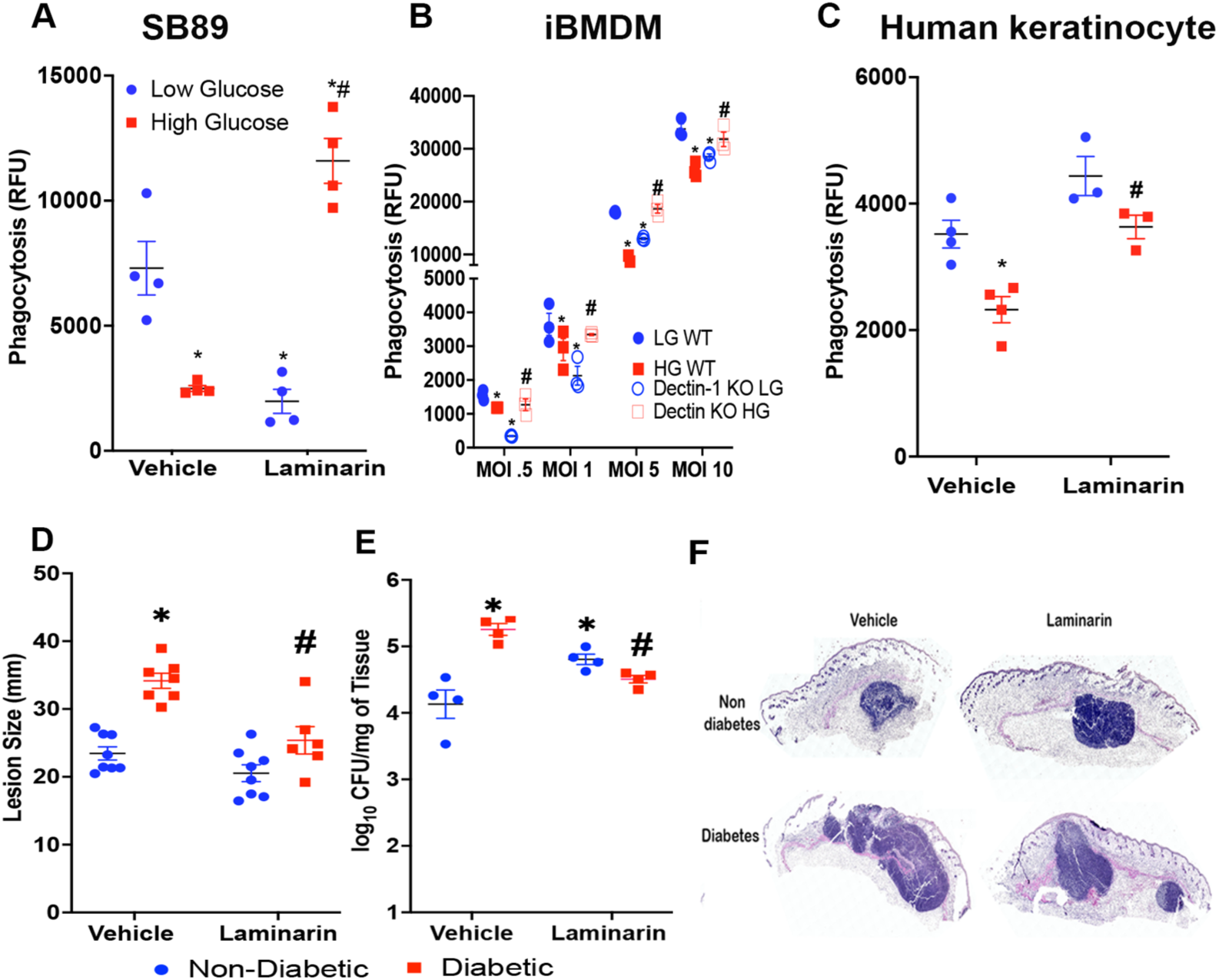
Dectin-1 inhibits *C. albicans* phagocytosis and is detrimental to host defense in diabetic mice. **A)** DMCLs (SB89)were cultured in low- or high-glucose conditions and incubated with laminarin for 30 minutes. Macrophages were infected with *C. albicans* for 60 minutes. **B)** iBMDs from WT and Dectin-1 KO mice were cultured in low- or high-glucose conditions and challenged with the indicated MOI for 60 minutes, followed by quantification of phagocytosis by RFU. **C)** Human keratinocytes immortalized were cultured in low- or high-glucose conditions and pretreated with laminarin for 30 minutes before C. albicans infection. **D-F)** C57BL/6 mice were rendered diabetic and treated with 250 mg/kg laminarin s.c. for 6 hours, followed by s.c. *C. albicans* infection. **D)** Lesion size, **E)** CFU and **F)** H&E staining were determined 24 hours after infection. Data represent mean ± SEM. *p < 0.05 vs non-diabetic or low-glucose control. # p<0.05 vs vehicle-treated non-diabetic mice or high-glucose-treated vehicle control by Mann–Whitney test. n = at least 4/group

To determine whether these events extend to human cells, we cultured keratinocytes (N/TERT) HK) in low- and high-glucose conditions and then challenged them with *C. albicans*. Our data show that human keratinocytes cultured under high-glucose conditions inhibit yeast ingestion, and that Dectin-1 antagonism restores phagocytosis in these cells (Figure 5C). These findings suggest that high-glucose conditions specifically alter Dectin-1 signaling, leading to inefficient ingestion of *C. albicans*.

Next, we investigated the mechanisms underlying Dectin-1-mediated inhibition of poor phagocytosis in cells exposed to high glucose. We first performed a phospho-kinase array on DMCLs cultured in low- and high-glucose media, then infected them with *C. albicans* for 60 min. We observed robust inhibition of phosphorylation of downstream Dectin-1 signaling molecules, including PLCγ, LCK, MSK-1, and CREB (47). We did not observe any difference between *C. albicans*-infected low- and high-glucose DMCLs for Lyn, STAT2, and 3 (Supplementary Figure 4). These data suggest that Dectin-1 inhibits the activation of enzymes involved in the phagocytosis machinery.

To evaluate the *in vivo* role of Dectin-1 in impaired host defense under hyperglycemic conditions, diabetic and non-diabetic mice were treated with laminarin (250 mg/kg, s.c.) for 6 hours prior to *C. albicans* infection. Laminarin treatment reduced lesion size and fungal burden (Figures 5D and 5E) in diabetic mice at day 1 post-infection. These data indicate that blocking Dectin-1 may be a therapeutic strategy to improve antifungal host defense in diabetic conditions.

### Dectin-1-mediated PGE_2_ inhibits phagocytosis in cells cultured in high glucose

Dectin-1 promotes PGE_2_ secretion in macrophages (21). Moreover, PGE_2_ suppresses macrophage antimicrobial functions (48). Here, we aimed to determine whether Dectin-1-driven PGE_2_ production impairs *C. albicans* uptake in diabetic conditions. Following *C. albicans* infection, we detected significantly higher levels of PGE_2_ in the skin of diabetic mice than in infected nondiabetic mice (Figure 6A). We also observed an increase in mPGES1 in the skin of nondiabetic controls (Figure 6A). This increase in PGE₂ was accompanied by increased mPGES-1 expression in diabetic skin 24 hours after infection (Figure 6B), consistent with increased prostaglandin biosynthesis.

**Figure 6.**
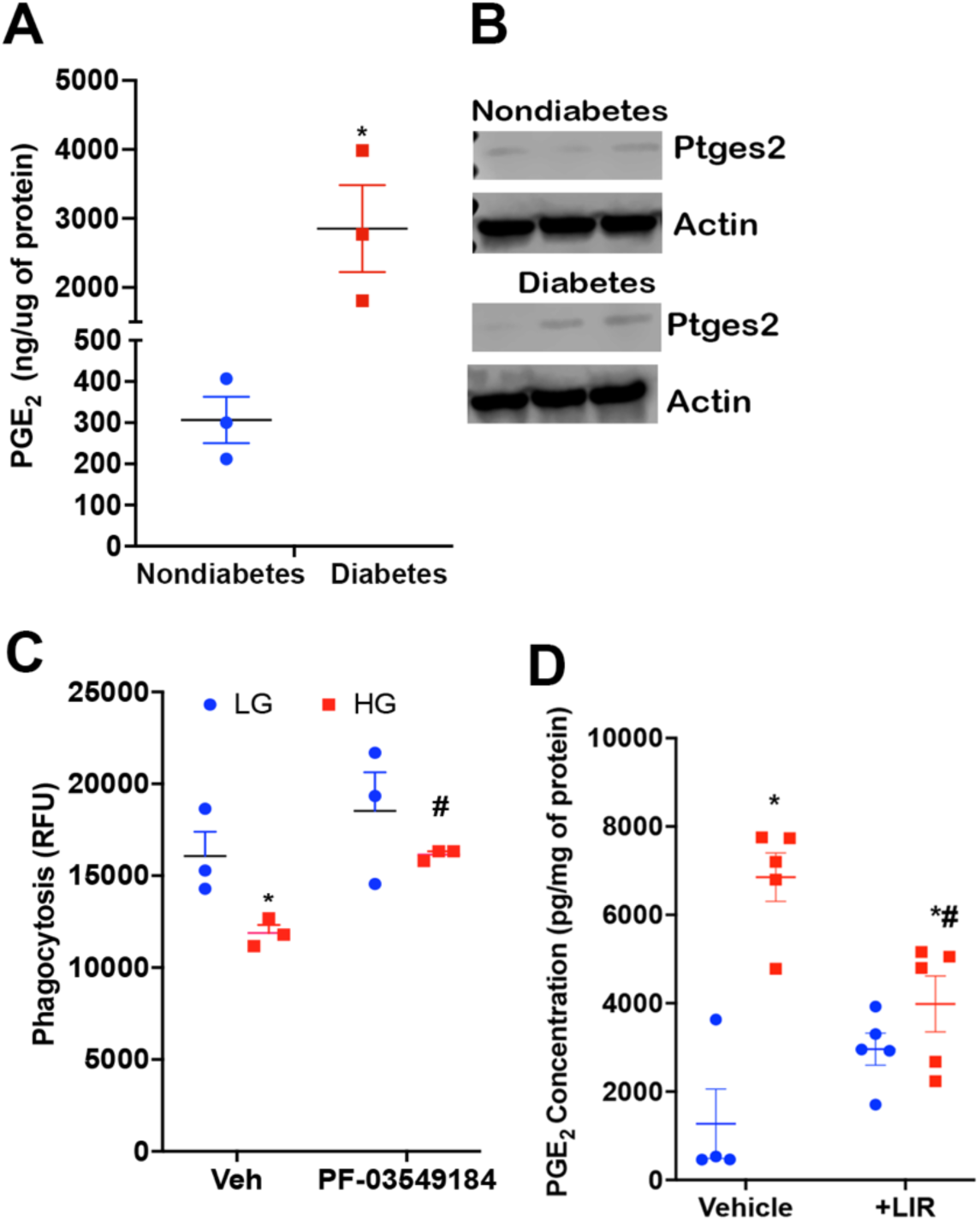
Increased PGE_2_ levels inhibit *C. albicans* phagocytosis. **A)** C57BL/6 diabetic and non-diabetic mice were infected with *C. albicans*, and PGE_2_ levels were measured in skin lysates as described in Methods. **B)** mPGES1 expression in the skin of infected diabetic and non-diabetic mice. **C)** DMCLs were cultured in low- (5 mM) or high- (25 mM) glucose and treated with the mPGES1 inhibitor PF03549184, 30 minutes before challenge with *C. albicans*. **D)** WT and Dectin-1 KO iBMDM were cultured in low- or high-glucose media, treated with PF03549184, and phagocytosis was determined as in **C**. **E**) Diabetic and non-diabetic mice were treated with liraglutide for 48 hours and infected as in **A**. Skin biopsies were isolated 24 hours after infection, and PGE_2_ levels were determined as in **A**. Data represent mean ± SEM. *p < 0.05 vs low glucose and #p<0.05 vs high glucose treated with vehicle control by Mann–Whitney test. n = at least 4/group

To determine whether elevated levels of PGE₂ were involved in low phagocytosis in high-glucose conditions, LG and HG macrophages (DMCLs and iBMDMs) were pretreated with PF-03549184, a selective mPGES-1 inhibitor, followed by challenge with *C. albicans.* Inhibition of mPGES-1 restored the phagocytic capacity of these macrophages compared with cells cultured in HG DMCLs, whereas little effect was observed under low-glucose conditions (Figure 6C). We observed the same results when WT iBMDMs were treated with an mPGES-1 inhibitor (Figure 6C). Next, we determined if Liraglutide decreases PGE_2_ abundance in infected diabetic and nondiabetic mice. Our data show that liraglutide prevents PGE_2_ generation in *C. albicans*-infected diabetic mice (Fig. 6D).

Overall, these data suggest that under high-glucose or diabetic conditions, Dectin-1 signaling promotes increased PGE_2_ production, which impairs macrophage phagocytosis, thereby worsening control of *C. albicans* infection. Reducing PGE2 levels, either by inhibiting Dectin-1 signaling or by restoring metabolic homeostasis, may attenuate inflammation and enhance microbial clearance in diabetic patients.

### PheWAS identifies associations between dysmetabolism and fungal infection within the CLEC7A SNP

To enhance the translational impact of this proposal, we examined whether Dectin-1 (*CLEC7A*) influences the development of fungal infections in individuals with metabolic dysfunction. We used a novel, unbiased genomic approach developed at Vanderbilt (49). Phenome-wide association studies (PheWAS) are valuable tools for identifying genetic variations in patient populations and linking genotypes to clinical outcomes(50) . We conducted a PheWAS analysis in a broad, disease-independent cohort of approximately 240,000 patients in BioVU (Vanderbilt’s biorepository of DNA from discarded blood collected during routine clinical testing, linked to de-identified medical records^)^ to explore disease associations with SNPs in genes involved in *CLEC7A* expression. BioVU offers a powerful bridge between mechanistic research and real-world clinical relevance, allowing us to assess a potential association of genetic variants to obesity, diabetes, and fungal infections. Based on the direction of the phenotype odds ratios (ORs), the results indicated that the *CLEC7A* SNP rs16910633 is associated with a higher likelihood of dermatomycosis (OR = 1.93, *P* = 0.003) and abnormal glucose (OR = 1.1, *P* = 0.01). These findings imply that *CLEC7A* is associated with an increased risk and severity of fungal infection in individuals with metabolic dysfunction.

## Discussion

Diabetes mellitus profoundly affects immune function, increasing susceptibility to and the severity of infections by impairing host defense mechanisms (51). It is well established that diabetes impairs microbial ingestion, killing, and chemotaxis in both mice and human cells (52). However, the intracellular events affected by high glucose remain elusive. In this study, we demonstrated that hyperglycemia inhibits cutaneous host defense against *C. albicans*, leading to increased fungal burden, exaggerated tissue damage, and defective pathogen containment. Using complementary *in vivo* and *in vitro* approaches, we identified an unexpected and detrimental role for the C-type lectin receptor Dectin-1 in shaping antifungal responses under hyperglycemic conditions. Rather than promoting effective fungal clearance, Dectin-1 signaling under high glucose suppresses phagocytic function and amplifies PGE₂ production, collectively contributing to inefficient fungal control. These findings reveal that metabolic stress rewires antifungal recognition pathways and uncover a mechanism by which diabetes predisposes to severe and persistent cutaneous candidiasis.

A key feature of diabetic skin infection in our model was the early inability to control fungal growth, as evidenced by a higher fungal burden within 24 hours. This was linked to heightened inflammatory infiltration, reflecting a dysregulated, overly aggressive yet ineffective inflammatory response (25). These findings support previous research showing that diabetes is not merely an immunosuppressed condition but involves maladaptive inflammation that damages tissue while failing to clear pathogens (9,41). Our results build on this concept by showing that, in fungal skin infections, early inflammatory signals are heightened in hyperglycemic skin, but these responses do not lead to effective antifungal action.

We and others have shown that skin infections in diabetes are driven by exaggerated inflammatory responses (9,25). Although we observed larger abscesses in *C. albicans*-infected diabetic mice, capsule thickness was similar between infected diabetic and nondiabetic mice (not shown). These findings contrast with our previous data from a murine model of MRSA skin infection in diabetic or obese conditions (38,39). In that model, infection in non-diabetic mice results in a well-delineated, compact abscess, whereas MRSA infection in diabetic or obese mice is characterized by a markedly thinner or poorly defined capsule (9). While the mechanistic basis for these differences was not investigated, we speculated that distinct microbial virulence programs may respond differently to hyperglycemic environments. Glucose availability has been shown to modulate complement evasion via the glucose-dependent expression of Hgt1 (53) and to reprogram transcriptional and stress adaptation pathways that enhance resistance to host-derived oxidative and cationic stresses at physiologically relevant concentrations (54)

Our studies are consistent with previous findings that diabetes is associated with increased abundance of microbes, including *C. albicans* (9,52,55–57). However, the mechanism underlying impaired ingestion of different pathogens remains to be determined. Here, we advance the field by showing that high glucose levels significantly impair fungal ingestion across multiple phagocytic cell types, including dermal, bone marrow-derived, and peritoneal macrophages from mice and human keratinocytes. This impairment affects both yeast and hyphal forms of *C. albicans*, indicating a broad defect in fungal uptake rather than a specific defect related to cell type or fungal morphology. Significantly, these results imply that hyperglycemia directly affects cell-intrinsic antifungal mechanisms in both immune and structural cells, potentially worsening tissue damage.

An important finding of this study is that Dectin-1, a receptor required for effective antifungal defense, adopts an inhibitory role under hyperglycemic conditions. Pharmacologic blockade of Dectin-1 with laminarin restored fungal ingestion in macrophages cultured in high-glucose media, and genetic deletion of *Clec7a* maintained phagocytosis despite metabolic stress. Moreover, *in vivo* laminarin treatment decreased lesion size and fungal burden in diabetic mice, highlighting a functional role for Dectin-1 signaling in poor infection outcomes. These findings challenge the conventional view of Dectin-1 as protective during fungal infections, suggesting that its signaling is highly context-dependent, especially in tissues under metabolic stress. The role of Dectin-1 in diabetes-related complications has indicated that elevated glucose levels increase Dectin-1 expression, which worsens insulin resistance, nephropathy, cardiomyopathy, and atherosclerosis across multiple experimental models (18,58–61).In these settings, genetic deletion of Dectin-1 in diabetic mice consistently reduced inflammation and conferred tissue protection. Additionally, high glucose increases Dectin-1 expression levels through a mechanism dependent on NF-κB activation (62). Although we did not directly investigate why Dectin-1 expression rises in macrophages and keratinocytes, previous studies have shown that AP-1 regulates Dectin-1expression via LTB_4_-mediated BLT1 activation (63). Whether this pathway is also affected under diabetic conditions remains to be determined.

Transcriptomic profiling of a newly generated dermal macrophage cell line revealed that exposure to a high-glucose environment induces a paradoxical state characterized by increased expression of fungal recognition receptors, including Dectin-1 and Dectin-2, and by suppression of downstream signaling components critical for effective antifungal responses, such as Syk, Malt1, and p38 MAPKs. Concomitant reductions in NF-κB pathway components indicate that hyperglycemia uncouples receptor engagement from productive intracellular signaling. Furthermore, phospho-array analysis showed that many kinases relevant to Dectin-1 signaling, including Src kinases, MSK1/2, and PLCγ, are affected by high glucose levels. These data reinforce the notion that although Dectin-1 expression is increased, intrinsic downstream pathways are negatively affected by hyperglycemia. This decoupling may result in sustained receptor activation in the absence of effective phagocytosis, promoting chronic inflammation while impairing pathogen clearance. Such a state mirrors features observed in other diabetes-associated inflammatory disorders, in which persistent pattern recognition receptor signaling drives tissue damage rather than resolution. Our lab is interested in determining how glucose negatively affects the activation of these molecules, not only in Dectin-1 signaling but also in other PRRs, including TLR2 and 4.

Mechanistically, we identified PGE₂ as a key downstream mediator linking Dectin-1 activation to impaired phagocytosis under hyperglycemic conditions. These findings are consistent with the well-established capacity of PGE₂ to suppress antimicrobial effector functions, including phagocytosis and reactive oxygen species production (24,64,65), while promoting vasodilation and leukocyte recruitment (65). We previously demonstrated that PGE₂ inhibits *Candida* ingestion by alveolar macrophages in a manner dependent on the cyclic AMP/PKA/PTEN axis (66). However, whether this signaling program is similarly affected in diabetic conditions remains under investigation.

Increased PGE_2_ production has been observed in various conditions associated with heightened susceptibility to infection, such as protein-calorie malnutrition, cancer, infancy, aging, cystic fibrosis, cigarette smoke exposure, bone marrow transplantation, and HIV infection (41,65). These findings align with our current data, which show that diabetic mice produce higher PGE_2_ levels than infected non-diabetic mice after *C albicans* skin infection. Interestingly, unlike our overall observations, MRSA infection in diabetic mice results in decreased PGE_2_ levels, and topical PGE_2_ analog misoprostol treatment restores host defense under diabetic conditions (25). Notably, the same enzyme is affected differently by multiple infectious agents: mPGES1 expression is suppressed in MRSA-infected diabetic mice, while Candida infection increases the expression in diabetic animals. These unexpected and intriguing findings suggest that a high-glucose microenvironment may not always enhance Dectin-1, and underscore that PGE_2_ changes are context-dependent, indicating that therapeutic strategies to boost or inhibit PGE_2_ should consider the specific pathogen involved.

Studies indicate that patients who achieve glycemic control through insulin therapy experience improved outcomes during infectious diseases (67,68), Building on this concept, over the past decades, a new class of therapeutics for glycemic control and weight management has emerged: GLP-1 receptor agonists. Currently, GLP-1–based therapies, such as semaglutide and tirzepatide, represent highly effective treatment options for individuals with type 2 diabetes and obesity, providing robust glycemic control and sustained weight loss, while also reducing cardiovascular and renal morbidity and mortality (69). Our data indicate that improving metabolic control contributes to the restoration of antifungal immunity. In diabetic mice, treatment with liraglutide, a GLP-1 receptor agonist, reduced lesion size, fungal load, and tissue damage, while also normalizing PGE₂ levels and partially rebalancing anti-inflammatory cytokine production. Collectively, these findings suggest that metabolic interventions can indirectly modulate innate immune signaling pathways, supporting the concept that host-directed therapies targeting metabolic inflammation may improve outcomes in individuals with diabetes.

Several limitations of this study should be recognized. Although pharmacological inhibition with laminarin and genetic deletion indicate a role for Dectin-1, additional studies are needed to clarify the specific downstream signaling pathways involved and the contributions of distinct PGE₂ receptor subtypes. Additionally, while the STZ model effectively mimics key aspects of hyperglycemia, future studies in obesity-related or type 2 diabetes models are necessary to assess whether these results are broadly applicable. Nevertheless, the combined evidence from *in vivo,* cellular, and transcriptional analyses strongly suggests that hyperglycemia alters antifungal recognition pathways, shifting them toward inflammatory but ineffective immune responses.

In summary, this study reframes susceptibility to fungal infection in diabetes as a consequence of dysregulated innate immune sensing rather than insufficient pathogen recognition. Hyperglycemia redirects Dectin-1 signaling from a protective antifungal pathway into a driver of inflammatory dysfunction via PGE₂-dependent suppression of phagocytosis. These findings represent a conceptual shift in our understanding of innate immunity in metabolic disease and suggest that targeting maladaptive sensing pathways, or their inflammatory downstream lipid, may offer new opportunities for host-directed therapies in individuals with diabetes.

## ACKNOWLEDGMENTS

We would like to thank the Serezani laboratory for their input and support. We also extend our thanks to Michail Lionakis (NIH) for providing the *C.albicans* strain SC5314. This work was supported by NIH grants 1R01AI180777-01, R01DK122147 (to CHS), 5R21AI171466 and 5R01AI168210 (to R.M.-B.), and HL174961 and NIH R01 HL159487 to AD. D.R was funded by 5T32AI112541-07 Genomic sequencing was performed in collaboration with the VANTAGE core services at Vanderbilt University, which is supported by the Vanderbilt Ingram Cancer Center (P30 CA068485) and the Vanderbilt Digestive Disease Research Center (P30 DK058404).

---

**Supplementary Figure 1.**
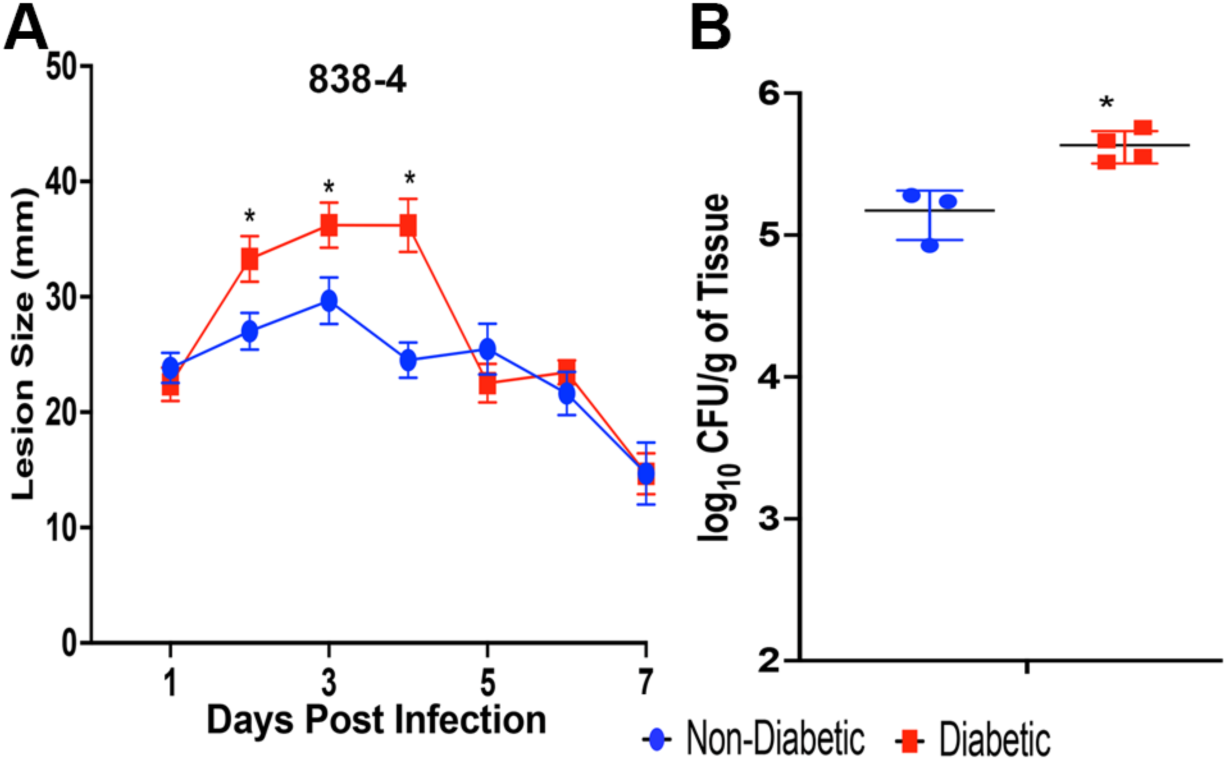
Diabetes enhances susceptibility to different strains of *C. albicans*. C57BL/6 mice were rendered diabetic for 30 days and infected s.c. with commensal *C. albicans* strain 838-4 **A)** Lesion size was determined daily for 7 days. **B)** CFU were determined in mice as in **A**, infected for 24 hours, and followed by biopsy collection. *p<0.05 vs non-diabetic. N=3-5/group.

**Supplementary Figure 2.**
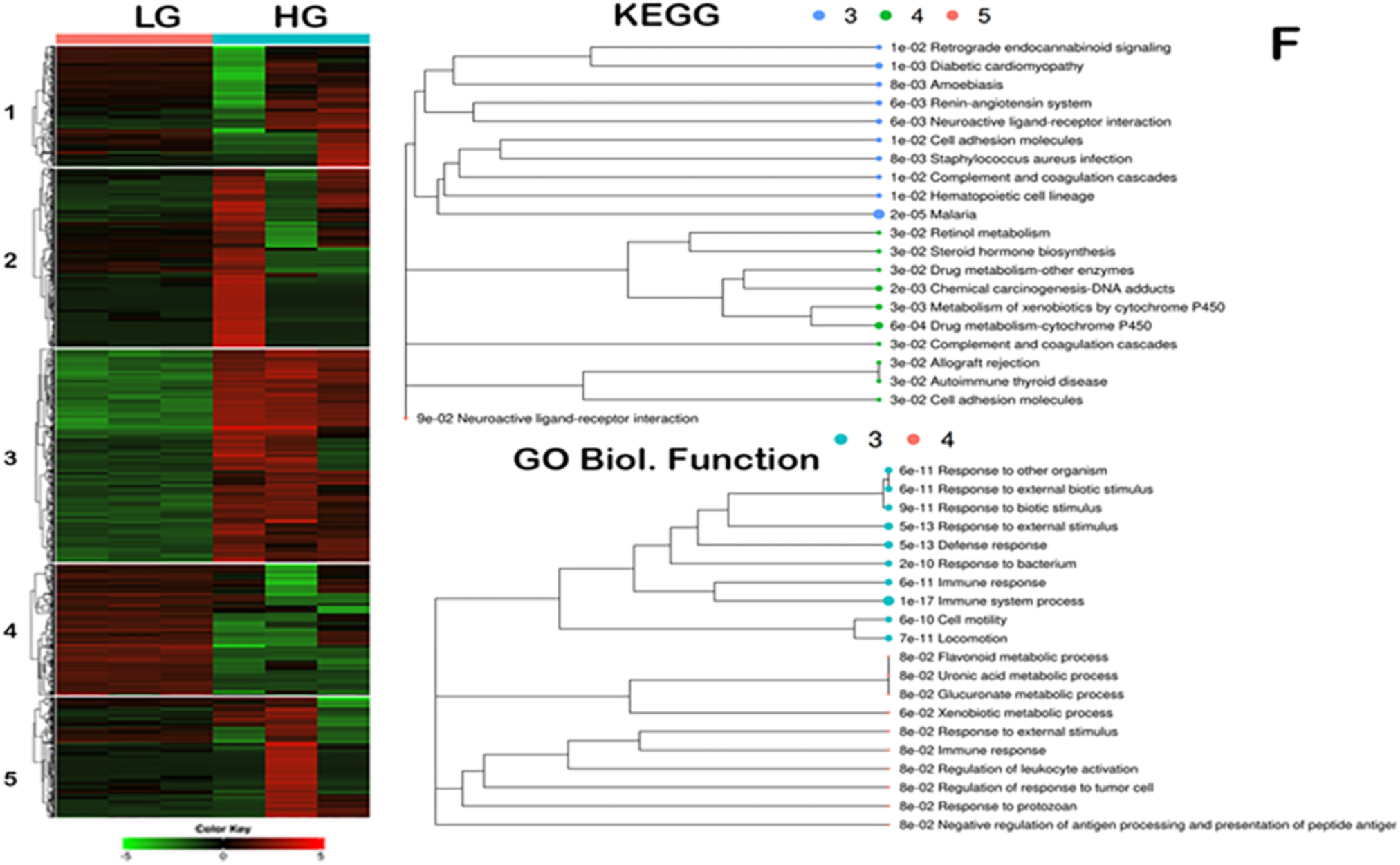
High glucose alters PRR expression in infected DMCLs. **A)** DMCLs were cultured in low- or high-glucose conditions and challenged with *C. albicans* for 12 hours. Heatmap and gene clustering of genes enriched in DMCLs cultured under low- or high-glucose conditions. The iDEP Analysis Toolkit and FDR < 0.05 were used to perform **(B)** KEGG enrichment and **(C)** Gene ontology – biological function of differentially clustered 3-5 derived from the genes clustered as in A. Data represent at least 3 independent experiments, and analysis was performed as described in Materials and Methods.

**Suppl. Fig. 3.**
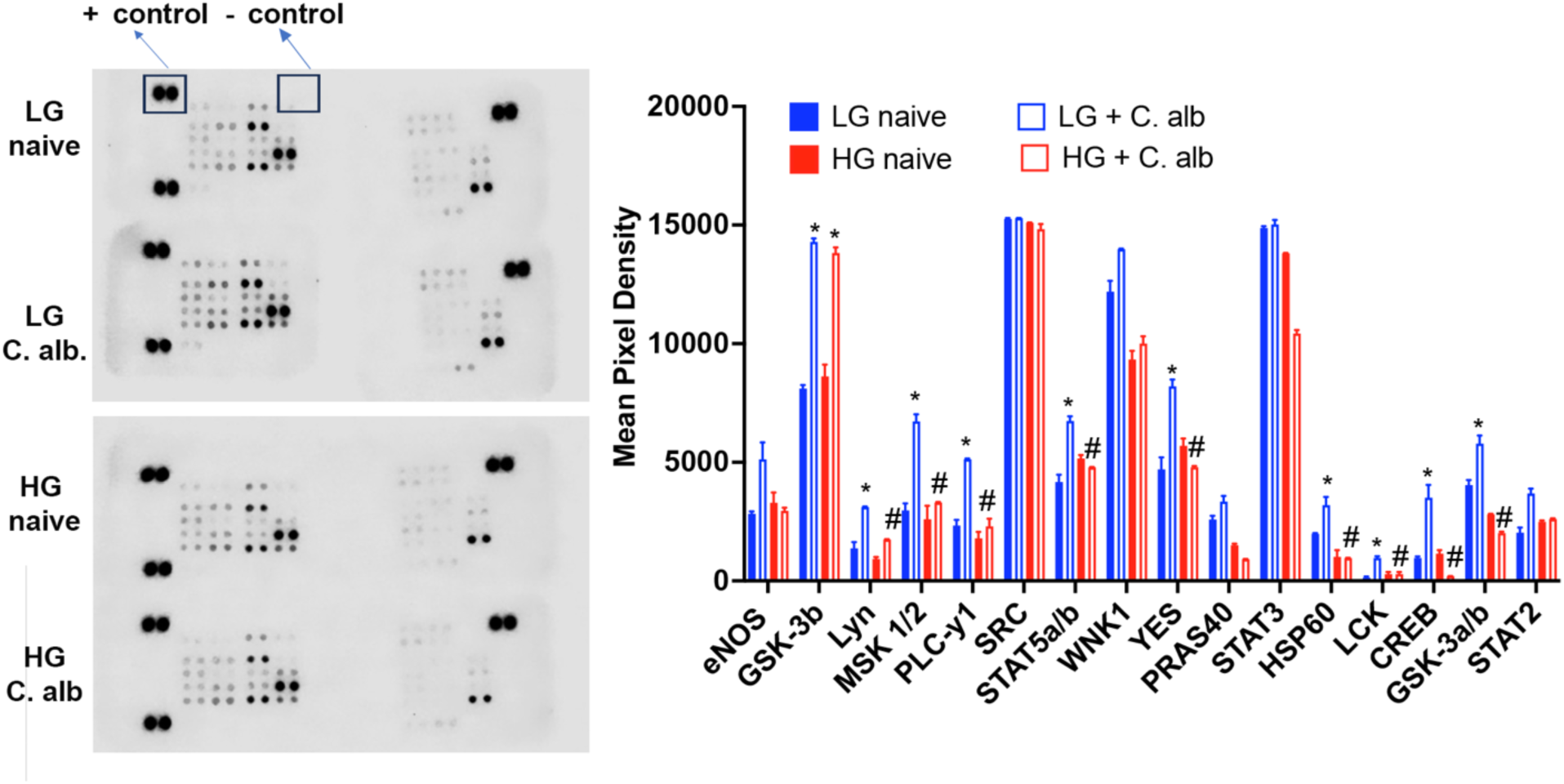
High glucose inhibits activation of dectin-1 downstream effectors. Left) DMCLs were cultured in low- or high-glucose medium and infected with *C. albicans* for 2 h. Cell lysates were prepared and subjected to phosphor-array as described in Methods. The image represents three independent experiments. Right) Quantification of spots/pixels from three independent experiments. Data represent mean ± SEM. p < 0.05 by Mann–Whitney test. n = 3/group

